# Differential patterns of cortical expansion in fetal and preterm brain development

**DOI:** 10.1101/2025.05.19.653958

**Authors:** Mariana da Silva, Kaili Liang, Kara E. Garcia, M. Jorge Cardoso, Emma C. Robinson

## Abstract

The human cerebral cortex undergoes a complex developmental process during gestation, characterised by rapid cortical expansion and gyrification. This study investigates in vivo cortical growth trajectories using longitudinal MRI data from fetal and preterm cohorts. We employed anatomically constrained multimodal surface matching (aMSM) to quantify cortical surface area expansion and compare *in-utero* versus *ex-utero* cortical growth using cortical surface data from 22 to 44 weeks post-menstrual age (PMA), acquired as part of the Developing Human Connectome Project (dHCP). Our findings revealed distinct regional and temporal growth patterns during the 2^nd^ and 3^rd^rd trimesters of healthy fetal cortical expansion. *Ex utero* brain development following preterm birth was shown to follow a modified trajectory compared to normal gestation, with potential implications for cortical organisation. Our methodology, combining biomechanically constrained surface registration with high quality fetal and neonatal imaging, provides a powerful framework for understanding early cortical development and deviations associated with preterm birth.

## Introduction

The human cerebral cortex undergoes a complex and lengthy development process that begins in the initial weeks of pregnancy and includes multiple overlapping mechanisms that occur at varying times and rates in different cortical areas (Budday et al., 2015; Dubois & Dehaene-Lambertz, 2015). Cortical development is complex and dynamic, requiring precise orchestration of events: starting with neurogenesis and proliferation from the sub-ventricular zones at the centre of the brain; followed by radial migration of neurons outwards to form the six layers of the cortical-plate, and post-migration mechanisms including dendritic arborisation (Budday et al., 2015; Kostović et al., 2019).

Folding allows for the cortex to attain a much greater surface area relative to the brain volume. While the mechanisms of cortical folding are not yet fully understood, growing evidence shows that they are a combination of genetic, cellular, and mechanical factors (Llinares-Benadero & Borrell, 2019; Striedter et al., 2015). Following initial formation of the insula (at around 14 weeks), more generalised patterns of cortical folding start at approximately 20 weeks, all driven by biomechanical stresses that result from the greater tangential expansion of the superficial cortical layers, relative to the tissues below (Garcia et al., n.d.; Richman et al., 1975).

In healthy development, gyrification develops according to a specific spatio-temporal schedule during normal gestation, with primary folds forming around 22 weeks of gestation, secondary forming from around 30 weeks, and tertiary folds continuing to evolve after birth. Primary folds are deeper and show greater consistency in location across different individuals; whereas, secondary and tertiary folding patterns show much greater variation, both in terms of location and orientation. Considerable evidence suggests that emergence and shape of primary folds are largely governed by genetic factors; however, the mechanisms that drive later folding patterns are less completely understood (Budday et al., 2015; Garcia, Kroenke, & Bayly, 2018; Greiner et al., 2021).

Classically, cortical folding has been studied through physics simulations that are most often tested in simplistic two-dimensional frameworks, representative of small sections of the cortex (Budday, Steinmann, & Kuhl, 2014; Holland et al., 2015; Tallinen et al., 2014). These models have informed understanding of the role that non-geometric factors, such as tissue properties, layer thickness, and growth rates (Budday, Steinmann, & Kuhl, 2014; Darayi et al., 2022), play in constraining biomechanical stresses during normal development. In Tallinen et al., 2016, similar models were scaled to simulations of whole brain growth, using a fetal brain at 22 weeks GA as a starting point for the simulations. However, while these showed clear replication of general trends of cortical folding, they could not completely replicate the folding patterns observed for that same brain at a later time.

To better understand the mechanisms that explain the heterogeneity of folding across individual brains, it is necessary to observe these changes in vivo and, in this regard, magnetic resonance imaging (MRI) - along with the availability of segmentation and cortical reconstruction algorithms - has emerged as a powerful tool. Past in-vivo studies of cortical development have largely relied on cross-sectional or longitudinal observations made from preterm neonatal scans (Dubois et al., 2008; Garcia, Robinson, et al., 2018; Moeskops et al., 2015). These align with postmortem observations (Z. Zhang et al., 2010), pointing to regional variations in cortical growth patterns that start in central areas before progressing from primary to association regions (Dubois et al., 2008; Garcia, Robinson, et al., 2018).

However, preterm infants are born at a time when many cortical developmental processes are still occurring, leaving them vulnerable to impaired cortical development. Cross-sectional studies, comparing term-born infants with very preterm (VPT) infants, suggest that VPT individuals exhibit reduced cortical folding at term-equivalent age within the insula, superior temporal sulci, and pre- and post-central sulci (Engelhardt et al., 2015), with this contributing to a reduced cortical surface area overall (Kline et al., 2020). These changes persist through childhood and adulthood (Kelly et al., 2023; Y. Zhang et al., 2015), with conflicting evidence as to whether cortical development in VPT individuals catches up or persists over time. Gorham et al., 2024 reported accelerated rates of cortical expansion between birth and late childhood (9/10 years) in frontal, temporal, and supramarginal/inferior parietal junction areas, but results from Kelly et al., 2024 indicate that cortical volumes of the temporal lobe remain significantly reduced in VPT individuals (at 7 and 13 years), with this largely being driven by reductions in cortical thickness. All of this points to widespread delays and alterations to whole brain development with Ball et al., 2014; Dimitrova et al., 2021 suggesting that this extends beyond morphological changes to also impact cortical microstructure and whole-brain connectivity.

Recent advancements in fetal MRI techniques now offer the ability to study the *in utero* development of cortical growth and gyrification across the second and third trimesters. Accompanied by increasing availability of open data collections such as Developing Human Connectome Project (dHCP) (Edwards et al., 2022) and a range of fetal atlases, this has supported the development of normative reference charts that summarise expected regional trends of cortical surface volume, area, curvature, and gyrification (Xu et al., 2022).

Yet regional analyses from cross-sectional populations may obscure subtle sources of individual variation and cannot inform the future design of tailored biomechanical models, capable of predicting growth trajectories in individual brains. In this study, our objective is therefore to compare *ex utero* patterns of cortical growth in preterm neonates with *in utero* patterns of cortical growth of fetuses, using longitudinal MRI data collected as part of the dHCP (Edwards et al., 2022). This builds from previous work by Garcia, Robinson, et al., 2018 that used biomechanically-constrained cortical surface registration (Robinson et al., 2018) to measure spatio-temporal patterns of whole-brain growth from longitudinal data of VPT neonates, scanned at fixed intervals (28 → 30, 30 → 34, 34 → 38). This study pioneered use of anatomical MSM (Robinson et al., 2018), which integrates a hyper-elastic biomechanical penalty on deformations of anatomical surfaces, as opposed to spherical surfaces, as done in the original MSM framework (Robinson et al., 2014). This enforces enforce smooth and biological-plausible deformations, that align with physics-based models (Bayly et al., 2013; Budday, Raybaud, & Kuhl, 2014; Garcia, Kroenke, & Bayly, 2018; Tallinen et al., 2014) and cross-sectional and regional measurements (Xu et al., 2022), while allowing for longitudinal patterns of cortical growth to be explored with much greater precision and detail.

Unlike Garcia, Robinson, et al., 2018, longitudinal scans from the dHCP were not acquired at fix intervals, but rather over an extended window with fetal scans acquired between 22 and 37 weeks gestational age (GA), health term born neonates scanned between 37 and 45 weeks, and preterm infants scanned shortly after birth (from 27 to 37 weeks) and near term equivalent age (37-45 weeks). Despite such confound variation, this paper shows that aMSM alignment captures expected patterns of cortical growth across second and third trimesters; with fetuses and preterms displaying differentiated patterns of cortical growth particularly within late developing areas - with fetuses displaying increased growth of the posterolateral parietal cortex, and preterms displaying accelerated relative growth in frontal and temporal poles.

## 2 Anatomical Multimodal Surface Matching (aMSM)

aMSM is an extension of the original multimodal surface matching (MSM) algorithm (Robinson et al., 2014) for spherical alignment of cortical surfaces. This works by first projecting cortical anatomies to a sphere using metric (i.e. areal and/or angle) preserving mappings; then an input (or moving) spherical mesh is deformed, until its cortical folds and/or patterns of areal organisation better overlap with that of a target (or reference) mesh (Glasser et al., 2016; Robinson et al., 2014, 2018). Deformations are constrained to remain biologically plausible through regularising displacements based on geodesic distances along the cortical sheet, rather than Euclidean distances in 3D (as would be applied during volumetric registration).

Spherical alignment is a commonly used approach for simplifying cortical mappings (Fischl et al., 1999; Yeo et al., 2009). The key difference between MSM relative to other spherical registration frameworks is therefore that MSM learns discrete rather than continuous displacement fields, in this way framing image registration as a label assignment problem, optimised as an Markov Random Field (MRF):

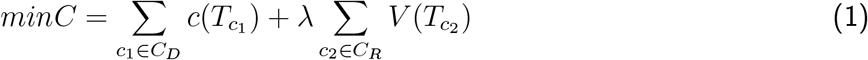

Here, deformations are constrained through warping a low-resolution, regular Control Point (CP) Grid, with labels representing a choice of possible deformations for each CP vertex (Fig 1c). The objective is to find the optimum set of labels that minimise a cost (1), which balances a data likelihood term (*c*(*T*_c*1*_)) with a regularisation penality (*V* (*T*_c*2*_)); where *c*(*T*_c*1*_) seeks to improve the overlap of cortical features (e.g. folds) between the moving and reference surfaces, and *V* (*T*_c*2*_) discourages biologically implausible deformation fields. This cost is calculated piecewise across cliques *c*_1_ and *c*_2_, with data terms most often estimated from unary cliques that consider the movement of individual CPs, and regularisation terms most often estimated from triplet cliques that penalise extreme distortions of triangular faces of the CP grid. *T* represents proposed affine transformations of each clique. This warp is then propagated to the (higher-resolution) input mesh using adaptive barycentric mesh interpolation (Robinson et al., 2018).

**Figure 1.**
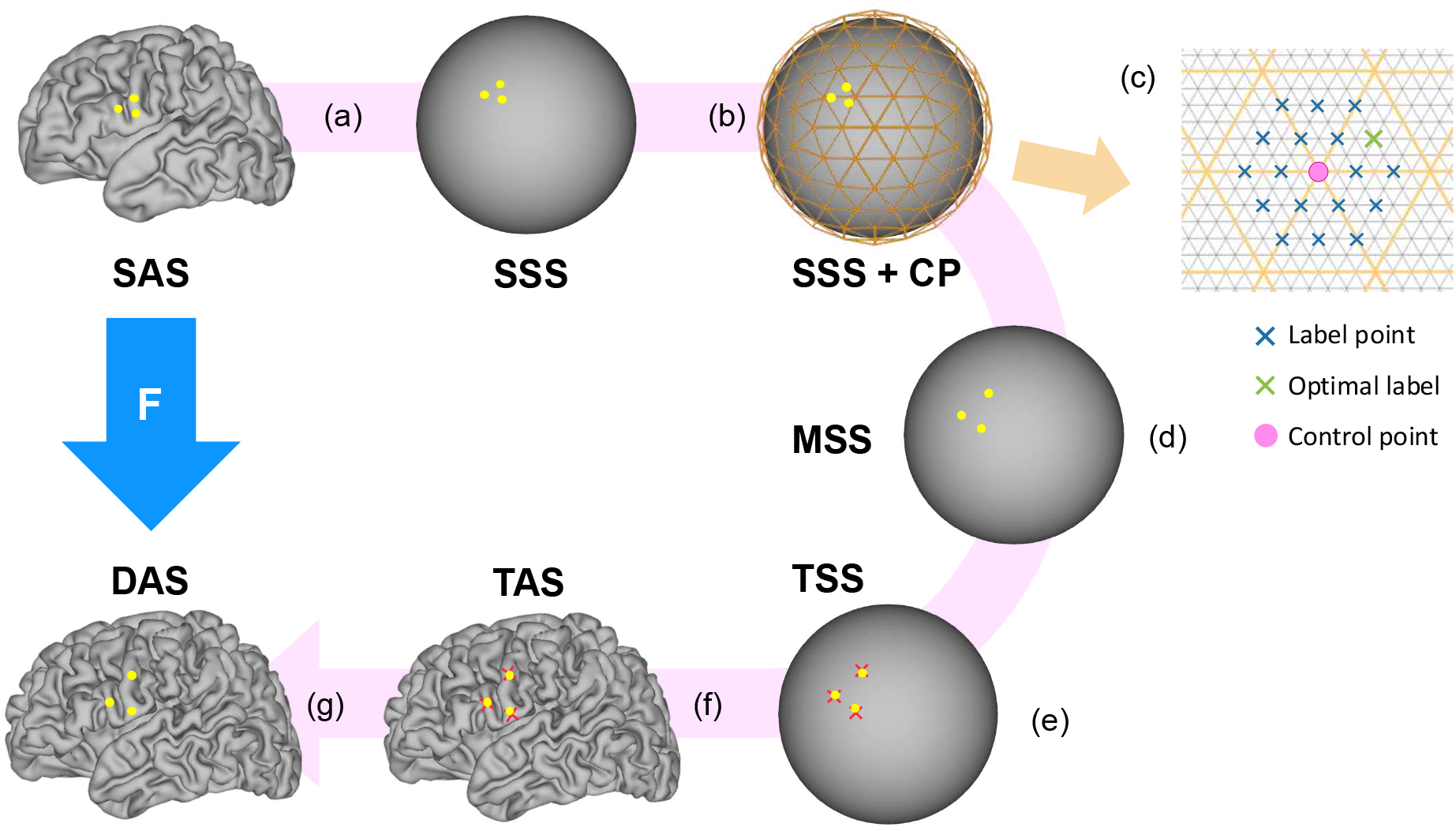
Overview of aMSM registration. a) The Source Sphere (SSS) and the Source Anatomical Surface (SAS) have point correspondence; b) Control-point grids (CP) guide the deformation of the SSS within a discrete optimization framework (c); d) The deformed spherical surface configuration (MSS) is derived from G using barycentric interpolation; e) Barycentric correspondences are established between vertices on MSS and TSS. f) The computed weights (from step e) are applied to corresponding points on the target anatomical surface (TAS). g) This results in a deformed anatomical surface (DAS), which retains the mesh topology of the source surface but adopts the shape of the target anatomical surface (Robinson et al., 2018).

In the most recent formulation of MSM, *V* (*R*_c*2*_) is imposed in the form of a hyperelastic strain energy function that penalises both areal and shape distortions. This is achieved using a variation of the Neo-Hookean equation, commonly used to describe the mechanical behaviour of soft biological tissues, including brain (Tallinen et al., 2014):

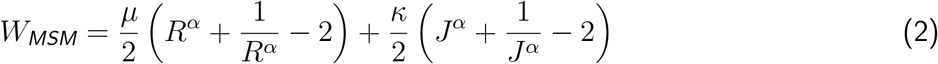

with *J* measuring areal distortions, and *R* measuring changes in shape (mathematical descriptions of these terms can be found in Robinson et al., 2018). Here, *µ* is the shear modulus, *κ* the bulk modulus and *α* is an integer ≥ 1. In spherical MSM this is imposed on distortions of the spherical mesh, but in aMSM this displacement is interpolated back to the anatomical mesh, thereby regularising deformations in the anatomical space instead. This makes use of the one-to-one vertex correspondence that exists between spherical and anatomical meshes, to infer the target locations of vertices on the anatomical moving mesh from their relative locations within triangles on the target mesh (Fig 1e-f). The work of Garcia, Robinson, et al., 2018 showed that biologically-informed choice of the parameters (*µ, »* and *k*) generates patterns of cortical expansion that are interpretable and align with expected trajectories of cortical growth. Comparing cortical expansion across subjects allows for the mapping of general trends of cortical growth. In this paper, we instead seek to use aMSM to compare trajectories of cortical growth between two cohorts that separately characterise *ex utero* and *in utero* cortical growth from 22-45 weeks GA.

## 3 Methods

### 3.1 Data

This study leverages longitudinal fetal MRI acquired from 72 fetuses and 90 preterm neonates, acquired as part of the dHCP. Fetuses were scanned *in utero* and then shortly after birth, preterm neonates were scanned shortly after birth and then near term-equivalent age. Patient demographics are shown in Table 1 and Fig. 2. We note that the age distribution for the first scan is highly scattered from 21 to 38 weeks GA, while the distribution for the follow-up scan is more concentrated over a range of approximately 5 weeks.

**Table 1.** Demographic information for studied cohorts: mean (range) or % (n).

| Variable | Fetal-neonatal, $n = 72$ | Preterm, $n = 90$ |
| --- | --- | --- |
| Gestational age at birth, wk | $40.2 \pm 1.3$ | $31.0 \pm 3.4$ |
| Birth weight, kg | $3.51 (2.54 - 4.34)$ | $1.54 (0.57 - 3.06)$ |
| Male | 51.39% (37) | 53.33% (48) |
| PMA at first scan, wk | $29.3 \pm 3.8$ | $33.5 \pm 2.4$ |
| PMA at final scan, wk | $42.2 \pm 1.4$ | $41.2 \pm 1.5$ |
| $\Delta t$ birth to first neonatal scan, wk | $2.4 \pm 0.9$ | $2.6 \pm 2.3$ |

**Figure 2.**
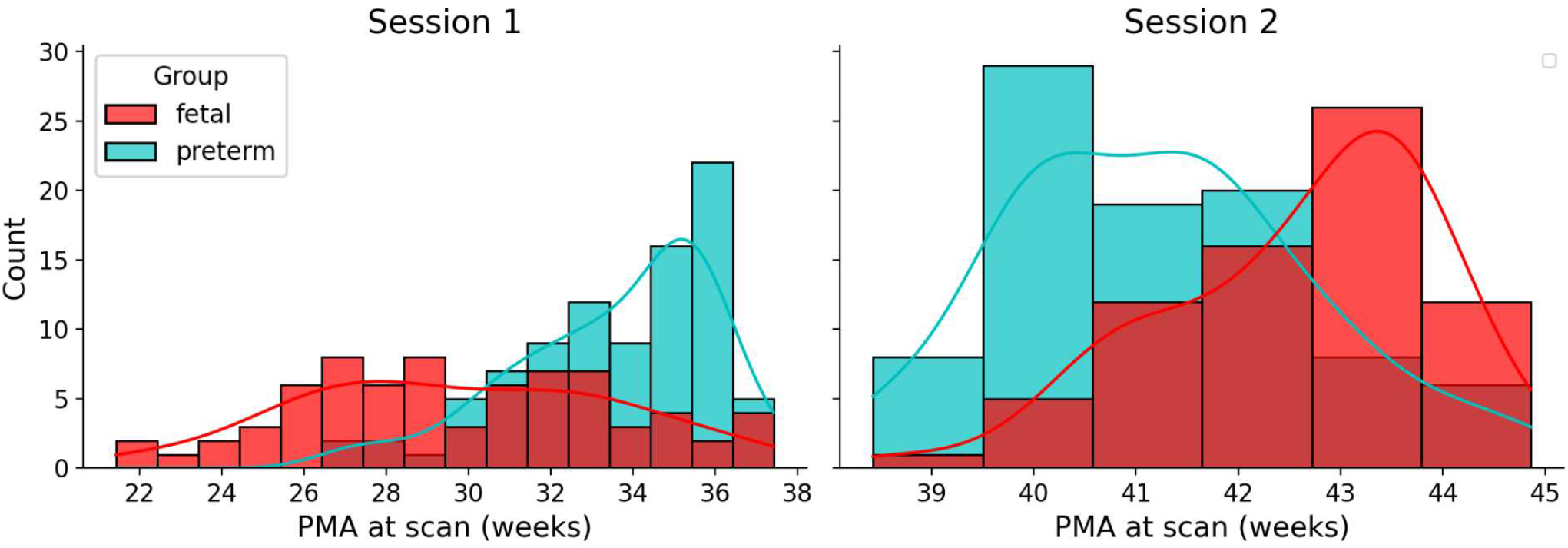
Histogram of age distributions at each scan for each of the groups.

#### Neonatal Acquisition and Pre-Processing

Neonatal imaging was performed using a 3T Philips Achieva scanner, equipped with a specialised neonatal imaging setup that involved a 32-channel phased array head coil. Anatomical imaging included fast spin-echo *T*_2_-weighted sequence, collected using a TR/TE = 12000/156ms, with in-plane resolution of 0.8mm × 0.8mm and slice thickness of 1.6mm, with an overlap of 0.8mm (0.74mm for *T*_1_-weighted sagittal). Scans were motion corrected and reconstructed to 0.8mm isotropic resolution; then passed through the dHCP structural pipeline (Makropoulos et al., 2018), which performed Draw-EM tissue segmentation (Makropoulos et al., 2014, 2016), followed by surface extraction (Schuh et al., 2017), inflation and spherical projection to obtain the inner and outer cortical (white and pial) surfaces, midthickness and inflated surfaces, and spheres, all with vertex correspondence. Full details of the dHCP image acquisition are described in Edwards et al., 2022, with reconstruction and preprocessing pipelines are described in Makropoulos et al., 2018.

#### Fetal Acquisition and Pre-Processing

For fetuses, *T*_2_-weighted images were collected on a Philips Achieva 3T scanner from 6 distinct orientations, using a high-zoom multiband (MB) single-shot TSE sequence with an in-plane resolution of 1.1 × 1.1, slice thickness of 2.2 mm, -1.1 mm gap and TE=250ms. Volumes were reconstructed to an isotropic resolution of 0.5mm with slice-to-volume registration (Kuklisova-Murgasova et al., 2012). Because of differences in acquisition, combined with rapid changes in brain size and shape over gestation, it was necessary to adapt the dHCP neonatal surface extraction pipeline for the fetal data. Surfaces used in this paper were generated from manually edited segmentations - output from an adapted version of the DRAW-EM tissue segmentation tool, which was modified to use population-average tissue priors generated specifically from the fetal cohort. The manually corrected segmentations were then processed through the dHCP neonatal surface reconstruction pipeline to generate white, pial, midthickness, inflated and spherical surfaces.

In both cohorts, cortical curvature was estimated from the mean of the principal curvatures extracted from a shape tensor fit at each WW vertex; however, cortical surface area was calculated by summing up the area of all triangles within the midthickness surface. Both fetal and neonatal surfaces were then resampled with adaptive barycentric interpolation Robinson et al., 2018 from their native meshes to a regular icosphere mesh of 40962 vertices, and rescaled to normalise the total surface area across all scans, prior to performing registration.

### 3.2 Longitudinal Surface Alignment

#### aMSM Implementation Details

Anatomically constrained multimodal surface matching (aMSM) (Garcia, Robinson, et al., 2018; Robinson et al., 2018) was employed to establish point-to-point correspondences between longitudinal pairs of anatomical cortical surfaces, extracted from the same subject at different time-points during gestation. The registration was driven by cortical curvature maps, with cortical anatomy represented by the mid-thickness surface.

Using the standard protocol for aMSM registration, mappings were initialised with rigid (rotational) alignment; then non-linear mappings were optimised in a coarse-to-fine fashion, using icospheric CP grids of increasing resolution (order) - starting with ico-2 (162 vertices), then ico-3 (642 vertices), and finally order 4 (2562 vertices); anatomical and image grid resolutions were set 2 orders higher at each iteration (4, 5, and 6) with ico-5 having 10242 vertices and ico-6 having 40962 vertices. Parametrising optimisation in this way helps to enforce smoother mappings through first aligning of coarse (or more prominent) shape features, before subsequently refining finer details.

For all experiments, the bulk modulus was defined as 10 times greater than the shear modulus based on previous experiments run in Garcia, Robinson, et al., 2018. However, λ (Eq 1) was tuned to optimise the trade off between feature matching and smoothness of the warp. For this, a subset of 10 subjects were selected, spanning an even distribution of GA at scan. aMSM alignments were then run for all examples, over a range of λ (from 1*e*^−5^ to 1). The optimal λ was then selected from plotting the feature correlation against the 95th percentile of the estimated deformation strain, with distortion and correlation increasingly linearly with decreasingly λ to a point, after which distortion increases exponentially for little or no similarity gain. We therefore select a λ at the top of the linear range.

For more details on the parameterisation and optimisation of aMSM please refer to the online documentation ^1^. Following alignments of all subjects, results were visually quality checked for goodness of alignment by comparing overlap of the curvature feature maps between the registered and reference surface. To avoid possible bias in the analysis related to the direction of registration, as done in Garcia, Robinson, et al., 2018, aMSM registration performed was performed in both directions, i.e. from the younger to the older surfaces (forward registration) and from the older to young (reverse registration). The warp from the reverse registration was then inverted and averaged. Note we perform surface rescaling prior to the registration (in order to minimize the effects of large overall brain size differences in the aMSM optimisation), but then applied the calculated warps to the original native surfaces (non-normalised) in order to estimate the final deformed surface.

#### Calculation of Surface Area Expansion

After obtaining vertex correspondence between the pairs of surfaces, the expansion of the cortical surface (*G*_*SA*_) was calculated as the areal distortion *J* for each triangle of the moving mesh (Fig. 3), comparing its initial configuration to its deformed configuration (output from aMSM); this corresponds to the areal difference between triangles:

**Figure 3.**
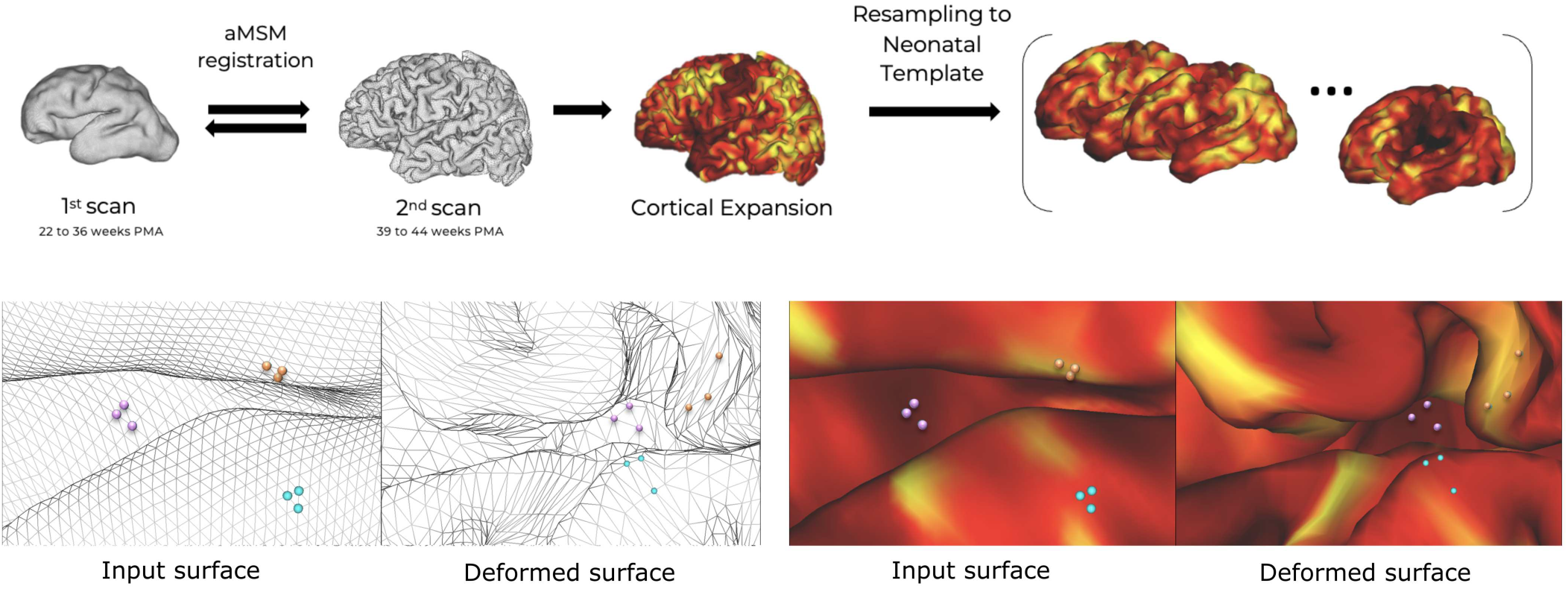
Overview of methodology for calculation of surface area growth maps. Top panel: aMSM registration was performed between younger and older scans in both directions, and averaged to create a mean registration warp and obtain point correspondence between vertices. Cortical expansion maps (*G*_*SA*_) were calculated from element-wise areal distortions between younger and corresponding registered surfaces. *G*_*SA*_ maps were smoothed, resampled to template space and downsampled. Bottom panel: Close-up on anatomical surface meshes showing correspondence of elements between input and deformed surface and corresponding vertex-wise growth. In the example, the orange triangle has higher *G*_*SA*_ (larger deformation) than the purple triangle.

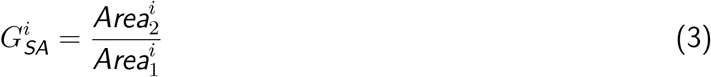

where *Area*_1_ is the area of triangle *i* in the younger surface and *Area*_2_ is the area of the corresponding triangle in the registered surface. These are averaged across the triangles of a vertex in order to compute vertex-wise values of *G*_*SA*_.

^1^Multimodal Surface Matching (MSM) Documentation

Relative growth maps, *rG*_*SA*_, were computed using the same method but for the results of the registration applied to the normalized surfaces (i.e. rescaled to the same total surface area).

### 3.3 Group Analysis

To compare population-level trends between preterm neonates and fetuses, all individual surfaces were aligned to a population-average 40-week GA neonatal surface template (Bozek et al., 2018; Williams et al., 2023). Noting that neonates and fetuses develop rapidly in size and shape, t’he dHCP generated a spatio-temporal surface template, which changes smoothly for each PMA. Aligning individual subjects into this space, required surface registration between that individual’s second scan and the template from its corresponding PMA i.e. for a subject whose second scan was acquired at 42 weeks and 3 days PMA, they would first be aligned to the 42 week surface template; this mapping would then be concatenated with pre-calculated template-to-template warps (from 42 → 41, then 42 → 41) in order to map that subject into the 40-week template space. All registrations were driven to align coarse scale patterns of cortical folding (sulcal depth maps) between source and template. Cortical growth maps were then projected through that mapping, and compared across and within groups using vertex-wise statistical permutation tests using FSL’s PALM (Winkler et al., 2014). To account for residual misalignments and reduce multiple comparisons, maps were downsampled onto a regular (ico-4) mesh of 2562 vertices and smoothed using a 4mm Gaussian smoothing kernel.

#### Statistical Models

PALM was configured to run General Linear Models (GLMs) which model cortical expansion for each vertex of the template. Considering an exponential model of growth, surface area was expected to increase as a function of age:

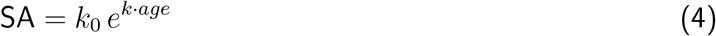

Allowing growth to then be modelled as:

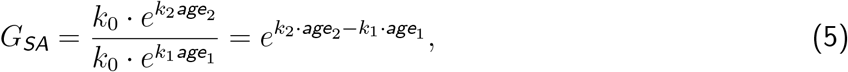

Which may be rewritten as a linear relationship through a ln (log_e_) transformation:

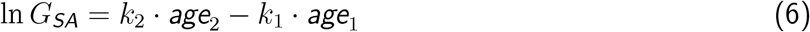

where *age*_2_ is the age at time-point 2 and *age*_1_ is the age at time-point 1.

If we assume that the exponential growth rate is constant (per vertex) for a fixed time-span, then the above equation can be re-written as:

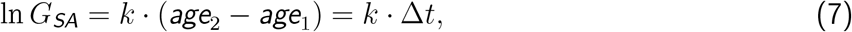

Noting that ln *G*_*SA*_ = 0 (equivalent to *G*_SA_ = 1) when the time-span between scans is 0, i.e. there will be no growth when *age*_1_ = *age*_2_.

We further consider the influence of sex on growth patterns, including it as a co-variate in our model. In order to account for small differences in age at the second scan, we also introduce the effect of *age*_scan2_ into our model.

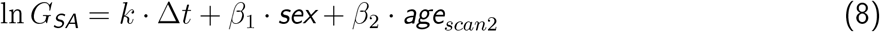

The the goal of the analysis is therefore to model the vertex-wise surface area growth rate, *k*, while deconfounding for the effects of the variables *sex* and *age*_*scan2*_ .

#### Temporal analysis of fetal growth patterns

In order evaluate temporal differences in fetal growth rates, we divide the analysis into two time-periods. The fetal-neonatal cohort was divided into two groups based on the age at *scan*_1_: 22 to 26 weeks PMA (corresponding to the end of the 2nd trimester) and 27 to 37 weeks PMA (3rd trimester). For this analysis, the (vertex-wise) GLM becomes:

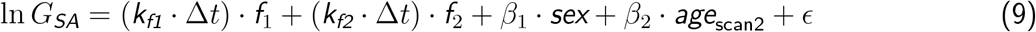

where *f*_1_ = 1, *f*_2_ = 0 for the subjects in the first group (younger fetal scan) and *f*_1_ = 0, *f*_2_ = 1 for the subjects in the second group. T-tests were performed to analyse if the growth rate (i.e. the slope between log *G*_*SA*_ and Δ*t*) is higher in the first time-period than in the second (*k*_*f1*_ >*k*_*f2*_) and vice-versa.

#### Comparisons between *in utero* and *ex utero* growth

No preterm scans were collected below 27 weeks PMA, therefore comparisons between *in utero* and *ex utero* growth was limited to group-wise comparisons between the older fetal-neonatal group (*f*_2_) and the preterm cohort, using the following (vertex-wise) GLM:

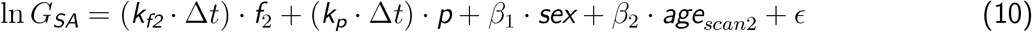

where *f*_2_ = 1, *p* = 0 for the subjects in the (older) fetal group and *f*_1_ = 0, *p* = 1 for the subjects in the preterm group. T-tests were again performed to compare the growth rates between groups.

For comparisons between relative growth maps (*rG*_SA_), analysis was performed to test whether the *means* of the two groups differ after adjusting for the confounding covariates.

PALM was applied using TFCE (Smith & Nichols, 2009) and 10,000 permutations for each contrast being evaluated.

#### Analysis of surface area at term-equivalent age

In order to compare surface area at term-equivalent age (corresponding to time-point 2 for both cohorts), vertex-wise surface area was calculated for the white matter surfaces registered to the 40-week neonatal template. Group comparison was performed using age, sex and total brain volume (calculated from the DRAW-EM segmentations in volume space) as variables.

## 4 Results

### 4.1 Global surface area development

We start by exploring global trends in surface area development for our entire cohort in a cross-sectional analysis (i.e using both scans from each subject as separate points). Fig. 4 shows the total surface area for each hemisphere, calculated from the native midthickness surface for each subject and each scan.

**Figure 4.**
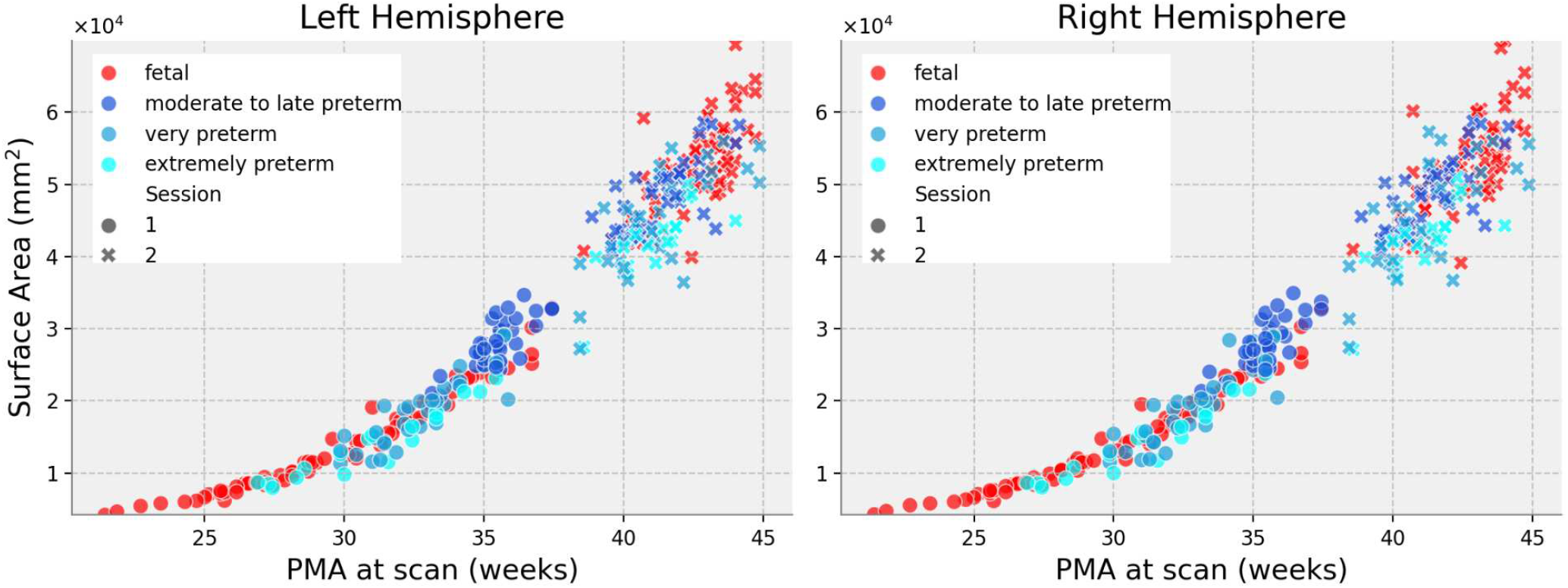
Total surface area (per hemisphere) calculated from the midthickness anatomical surfaces. We plot total area in both scans for each subject (scan 1, dots; scan 2, crosses). The preterm group is colour coded by prematurity degree (extremely preterm: GA birth *<* 28 weeks; very preterm: 28 weeks *<* GA birth *<* 32 weeks; moderate to late preterm: 32 weeks *<* GA birth *<* 37 weeks). In the subsequent analysis, we fit a single curve for the entire preterm cohort.

We note that global surface expansion follows an exponential trend in both the fetal group (*R*^2^ = 0.9736 and *R*^2^ = 0.9710 for left and right hemispheres respectively) and the preterm group (*R*^2^ = 0.9309 and *R*^2^ = 0.9258 for left and right hemispheres respectively). The global expansion rate was shown to be higher for the fetal group (*p <* 0.05) however the variance of the regression residuals is much larger for the preterm group (*p <* 0.0001).

### 4.2 Qualitative analysis of individual growth patterns

Fig. 5 shows results of individual cortical growth maps calculated from intra-subject longitudinal registration visualised in both young and older mid-thickness surfaces. These results highlight the rapid development of overall brain volume and surface area that occurs from 22 weeks GA to term and show that patterns of cortical expansion move from the central sulcus and Sylvian fissure to become more scattered around the frontal and parietal lobes.

**Figure 5.**
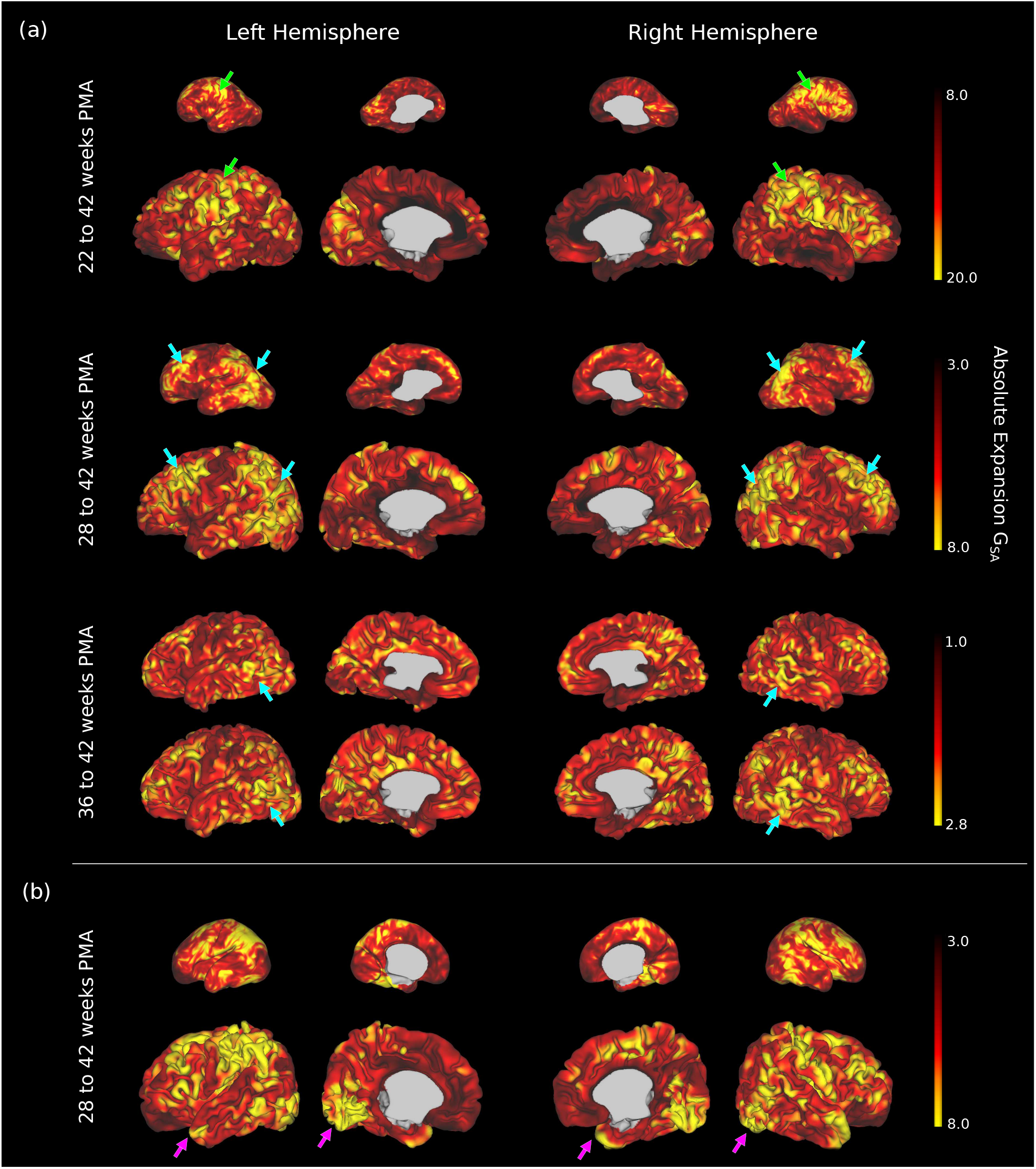
Absolute cortical surface expansion maps (*G*_*SA*_) for 4 individual pairs of surfaces, visualised in both the first and second scan. (a) Maps for 3 subjects born full term, scanned respectively at 22, 28 and 36 weeks in utero, and at 42 weeks. Between 22 and 42 weeks, the overall area growth is higher in the areas around the central sulcus and the Sylvian Fissure (row 1, green arrows). For growth maps covering only the 3rd trimester (rows 2 and 3) the patterns of expansion are higher in the lateral frontal and posterior parietal regions (blue arrows). (b) *G*_*SA*_ for a subject born prematurely (GA at birth = 24 weeks)

### 4.3 Cortical growth patterns in fetal development

Investigation of population-average trends across the fetal group align with the individual trajectories shown in Fig. 6, showing significant regional differences in growth patterns over the course of development, with growth in surface area higher for the final weeks of the 2^nd^ trimester relative to the 3^rd^ trimester for regions around the central sulcus and the insular region, coinciding with the emergence of primary and secondary folding in this period (*p <* 0.05). This corroborates the findings of Garcia, Robinson, et al., 2018, which also pointed to relatively increased growth of the sensory and motor regions over this period, with associative areas, such as the prefrontal cortex, expanding faster later in development.

**Figure 6.**
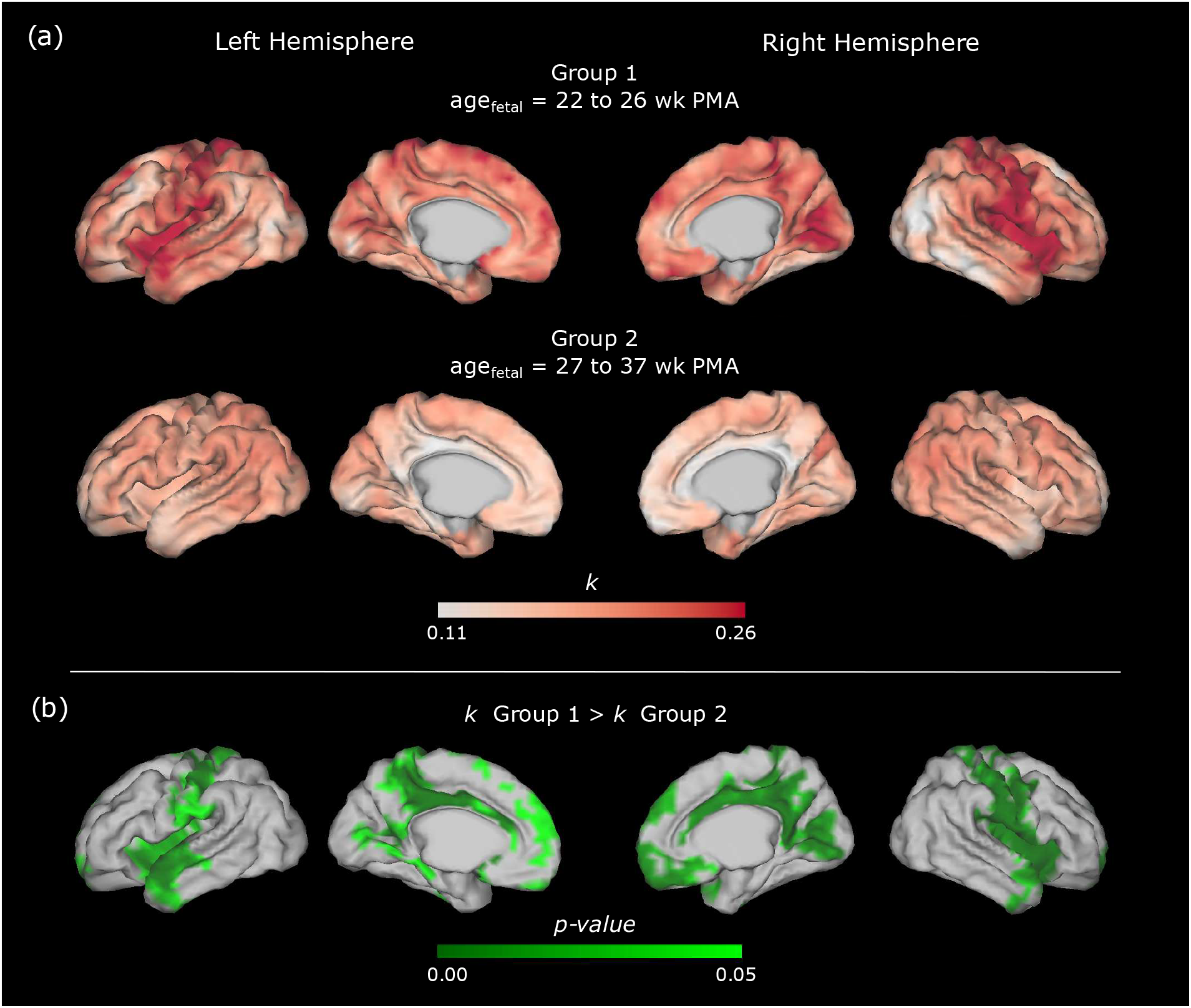
Analysis of fetal growth rates. (a) Vertex-wise *k* values (growth rates) for the period of 22 to 26 weeks PMA and 27 to 37 weeks PMA. (b) Growth rates are higher for the period of 22 to 26 weeks PMA for regions around the central sulcus, cingulate gyrus and insula; the opposite comparison (*k* Group 2 *> k* Group 1) did not show statistically significant differences.

### 4.4 Preterm growth patterns

Comparisons between *in utero* and *ex utero* growth were performed through group-wise comparisons between the older fetal-neonatal group (*age*_1_ *>* 27 weeks PMA, n = 52) and the preterm cohort (n = 90). We start by comparing the vertex-wise cortical area (calculated from the white matter surfaces after registration to a template space) between groups at term-equivalent age, corresponding to the second scan for all subjects (Fig. 7). When controlling for age at scan, sex and total brain volume, vertex-wise surface area was shown to be higher at term-equivalent age for the neonates born full-term, with this being statistically higher in regions around the frontal lobe, insula, and temporal lobes. Comparisons at time-point one were not performed since the distribution of ages is much larger for the first scan, and we expect surface area to highly change with age, not allowing for an accurate group comparison in a single template.

**Figure 7.**
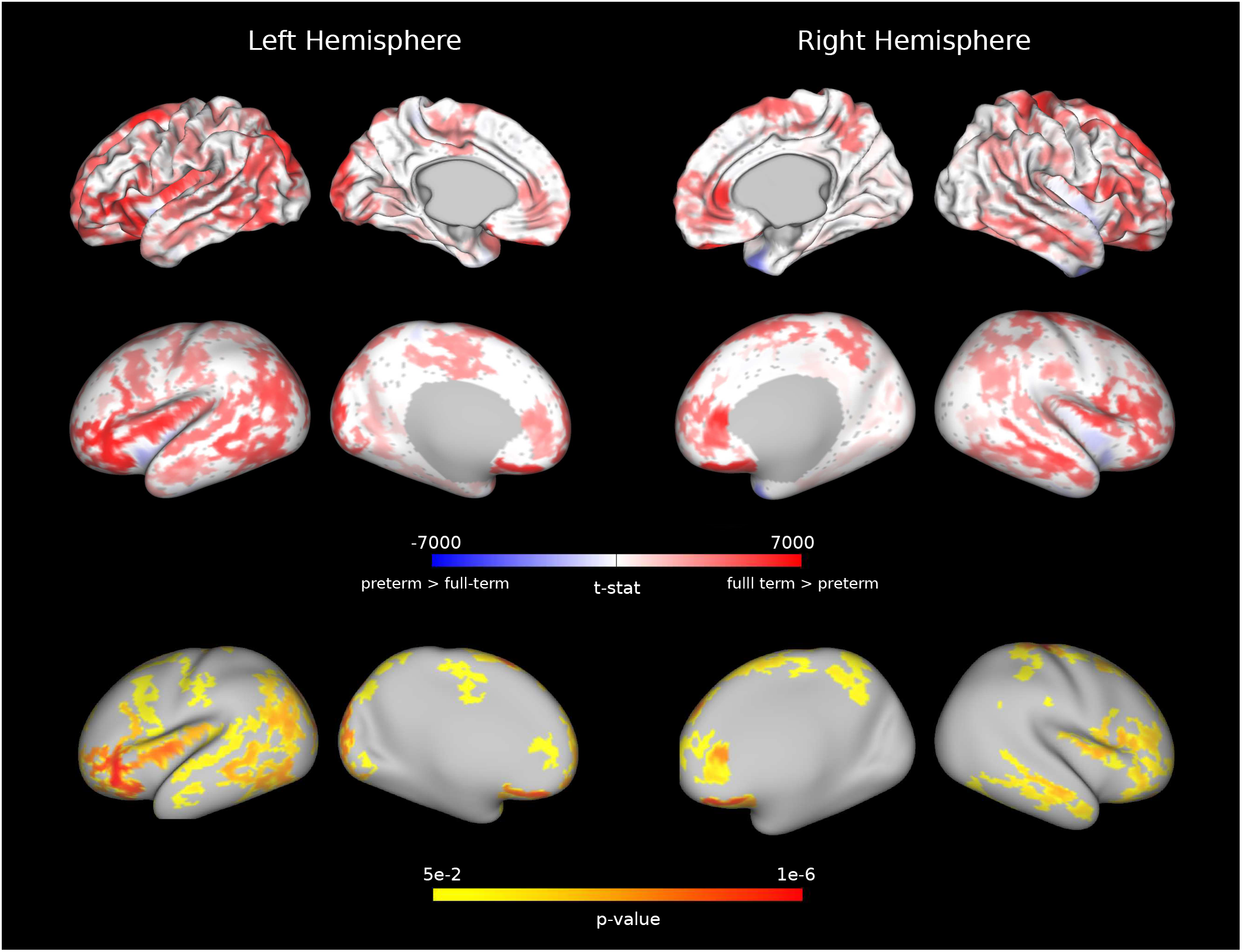
Results of group comparison (full-term vs preterm) for surface area at scan 2. T-statistics are shown on both the white matter and inflated surface for better visualisation. Regions of statistical significance (p *<* 0.05) are shown in the bottom panel, for *SA*_full-term_ *> SA*_preterm_. The opposite comparison did not show statistically significant differences.

Comparisons between absolute growth rates, *k*, in fetal and preterm development are shown in Fig 8; however, these did not survive multiple-comparison correction. It is possible that absolute growth patterns fail to reach significance due to confounding effects of overall surface area change, that change dramatically with age.

**Figure 8.**
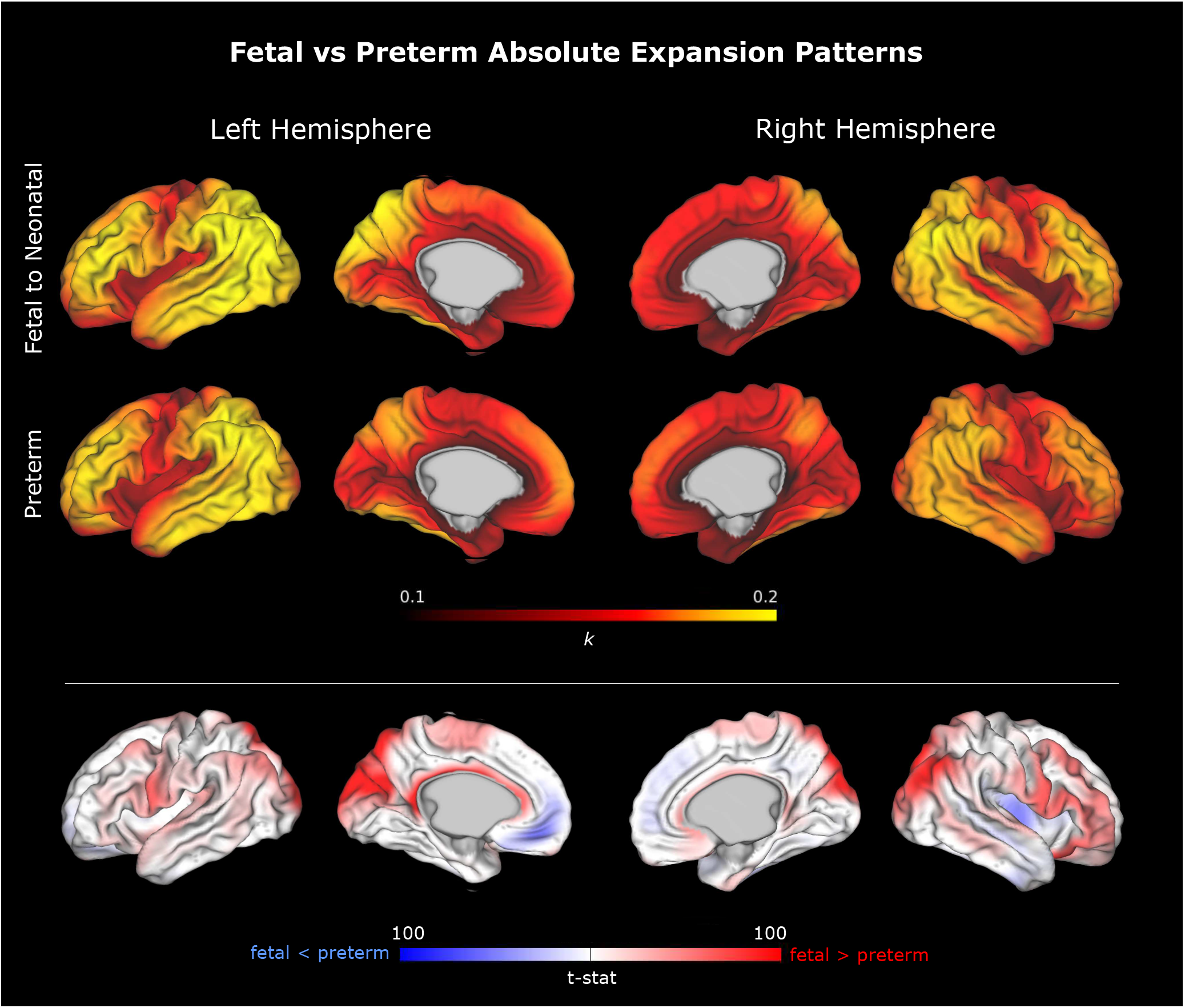
Vertex-wise *k* values (growth rates) for fetal and preterm brain development during the third trimester (27 to 37 weeks GA). The bottom row shows the results of group comparison (T-statistics), showing the fetal growth rates are higher in the areas around the central sulcus, parietal and occipital lobes (indicated in red), while preterm growth rates are higher in the insular region. However, this comparison did not result in statistical significance.

We therefore also compared relative growth after rescaling all surfaces to have the same total surface area, with the resulting maps highlighting which cortical areas were growing more or less than the population average across the entire cortex. Results in Fig 9 reveal that preterm neonates exhibited higher relative expansion in the temporal and frontal poles and insula, while fetal relative expansion was higher for the posterolateral parietal cortex. Interestingly, areas where relative growth is higher for the preterm group, such as the insula and frontal pole, correspond to areas where the surface area is lower at term-equivalent age as identified in Fig. 7, suggesting that preterm cortical growth is catching up with but not yet reaching the trajectory of the term-born cohort. These results align with Garcia, Robinson, et al., 2018, which measured relative cortical growth from a longitudinal preterm cohort, finding faster rates of growth in the lateral parietal, temporal, and occipital lobes, relative to central sulcus and insula, over later stages of preterm development (34 to 38 weeks), where this more closely matches the time-period analysed in this study.

**Figure 9.**
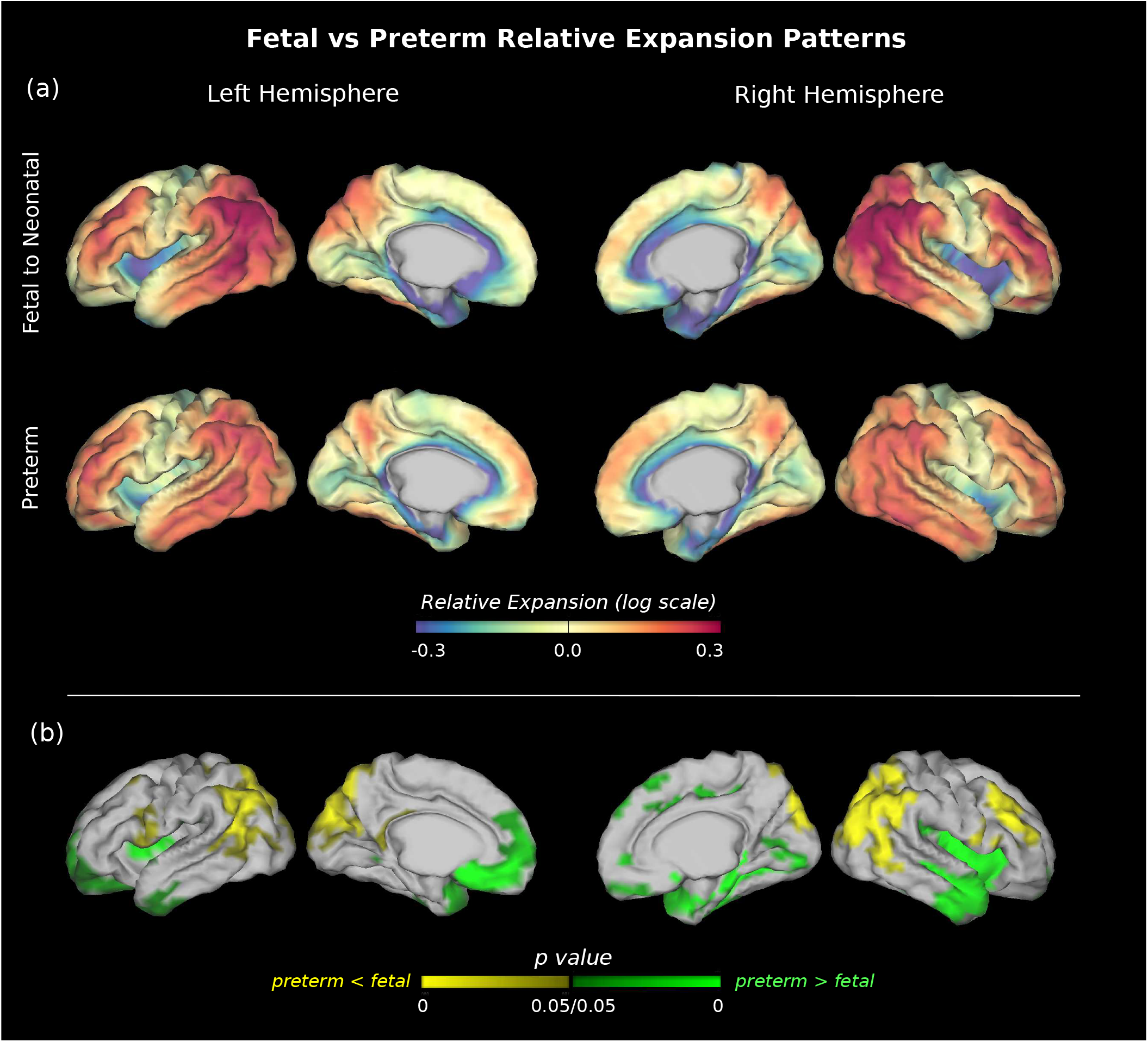
(a) Comparison of average relative cortical expansion (normalised for global brain expansion) for *in utero* and *ex utero* growth during the third trimester (27 to 40 weeks GA). (b) Group-wise statistical comparison reveals preterm neonates exhibit higher relative expansion in the temporal and frontal poles and insula, while fetal relative expansion was higher for the posterolateral parietal cortex.

## 5 Discussion and Future Work

In this work, we analysed cortical growth patterns in fetal and preterm brain development from longitudinal surface registration constrained by biomechanical equations. This framework allowed us to obtain point correspondence between scans of the same subject to calculate vertex-wise cortical growth patterns. By using a biomechanical regularisation within the registration model, we were able to estimate biologically plausible deformation warps according to the mechanical properties of the tissues.

The analysis of regional growth from longitudinal registration was able to identify patterns not captured by either total surface analysis or ROI studies. This can be seen when comparing the results from the PALM analysis on the fetal dataset - which showed growth rates vary not only regionally but temporally - with the initial analysis performed on total hemisphere surface area, which did not identify these differences. In previous studies (Xu et al., 2022) a single exponential curve for each ROI was used to cover the entire period of fetal development from 24 to 38 weeks. This can be justified from the fact that areas of larger ROIs can average out the effect of more subtle regional differences; particularly, the use of ROIs that split the regions adjacent to the central sulcus into the frontal and parietal lobes, as well as the regions surrounding the insula into the temporal and frontal lobes. By being able to measure vertex-wise growth without the constraint of pre-defined regions, we were able to identify regional differences in growth rates which agree with the literature. The fetal cortical gyrification (and therefore higher growth) has been showed to develop from highly conserved primary regions such as the insula and central sulcus, towards higher association areas later in development (Budday et al., 2015; Greiner et al., 2021). Therefore growth rates in surface area match the evolution of cortical folding from primary folds in earlier development such as the central sulcus, Sylvian fissure and cingulate sulcus, towards secondary and tertiary folding across the different lobes.

Healthy *in utero* and preterm cortical expansion patterns for the same period were compared through group analysis, showing distinct cortical growth patterns which are particularly highlighted when comparing *relative* growth patterns instead of *absolute* growth rates, which can be confounded by group differences in overall surface area at both time-points. Interestingly, as expected, differences between groups are more significant in regions that have previously been shown to develop at later stages of gestation. More surprisingly, some of the areas where relative surface expansion is higher in the preterm cohort - such as the frontal and temporal poles and insula - actually correspond to regions where the area is lower at term equivalent age, meaning that they are still underdeveloped. These results are in accordance with recent literature (Gorham et al., 2024) that suggests this differential development in the frontal and temporal lobes continues after birth and up until childhood.

This study however presents multiple limitations to the analysis. First, as visualised in Fig. 2, and as discussed in the introduction, the dHCP longitudinal cohort has a heterogeneous distribution of ages and time spans between scans. Secondly, only two time points are available for each subject, limiting the longitudinal analysis of growth to a single estimation of vertex-wise growth per subject, with the second time-point always corresponding to term-equivalent age. Due to this, growth estimates always correspond to total growth between the first scan, which has a high variation of ages (22 to 38 weeks PMA), and the scan at term equivalent age, with no intermediate scans. These limitations are reflected in the design of our group comparisons using GLMs: group comparison between younger and older fetal periods of development was performed considering a single rate of growth, *k*, (per vertex) for each of the groups; the same is the case for preterm and *in utero* development, where *log*(*G*_*SA*_) was consider to be linearly correlated to age. Future work should therefore look into modelling growth using a more complex regression analysis considering different growth rates *k* for smaller periods of development. This can be more easily achievable with multiple scans per subject (with a lower time-span between acquisitions), allowing for the estimation of more accurate weekly growth rates and the study of their evolution with time, as done in Garcia, Robinson, et al., 2018.

Another limitation of the present study, particularly affecting the comparison between fetal and preterm growth, are the differences in acquisition for fetal and neonatal MRI. While a dedicated processing framework was developed for the fetal dHCP cohort, including manual editing of surfaces in order to harmonise these with the dHCP neonatal pipeline results, acquisition differences can be reflected in systematic differences in metrics extracted from these surfaces, and future work should evaluate these differences.

As fetal MRI becomes increasingly more accessible and of higher resolution - together with the improvement in dedicated image reconstruction tools - further studies will benefit from larger longitudinal cohorts of healthy subjects, which will allow the construction of normative models of healthy fetal brain development. Cortical malformations could then be evaluated as deviations from these models. Future work should also consider incorporating other cortical development metrics into the analysis. For example, cortical thickness is known to evolve differentially during fetal development (Xu et al., 2022), and deviations in cortical thickness have been shown to affect cortical gyrification patterns (Budday, Raybaud, & Kuhl, 2014).

In conclusion, we were able to estimate detailed maps of cortical expansion using biomechanically-constrained longitudinal registration. These maps can be used to inform future models of cortical folding that more realistically consider heterogeneous growth rates across the brain surface. This analysis identified population patterns of healthy cortical development consistent with the literature but not observable in whole-brain or ROI-based MRI studies, and different them from *ex utero* development following preterm birth.

## Ethics

Data from the developing Human Connectome Project (dHCP) was approved by the United Kingdom Health Research Authority and informed written consent was given by the parents of all participants. This study followed the dHCP Open Access Data Use Terms and complied with institutional rules and regulations.

## Data and Code Availability

Data from the developing Human Connectome Project (Edwards et al., 2022) is used in this paper. The full dataset and documentation can be downloaded from The National Institute of Mental Health Data Archive (https://nda.nih.gov/editcollection.html?id=3955). Code used for data analysis is available at github.com/marianaftsilva/cortical-growth.

## Author Contributions

M.d.S: Conceptualisation, Formal Analysis, Investigation, Visualisation and Writing – Original Draft Preparation. K.L.: Formal Analysis, Investigation, Visualisation. K.E.G: Methodology, Conceptualisation, Writing-Reviewing and Editing. M.J.C.: Supervision. E.C.R.: Conceptualisation, Methodology, Supervision, Writing-Reviewing and Editing.

